# Quantitative trait loci mapping identifies genetic loci modifying MafA^S64F^-mediated dysglycemia

**DOI:** 10.64898/2026.11.06.764887

**Authors:** Mallory Maurer, Lizabeth Johnson, Dongsoo Lee, Brian S. Yandell, Mark P. Keller, Jeeyeon Cha

## Abstract

Increasing evidence has shown the role of genetic background on the penetrance of monogenic diabetes. A missense mutation of the pancreatic β-cell-enriched transcription factor MafA co-segregates with diabetes (MafA-MODY) and non-syndromic insulinomatosis. We recently investigated the effects of genetic background on MafA^S64F^-driven dysglycemia in genetically diverse mice harboring this mutation. Here we leveraged the effects of genetic background on phenotypic penetrance to map quantitative trait loci (QTL) potentially modifying glycemia in response to this variant. We investigated *Mafa*^S64F/+^ mice from an F2 intercross of the C57BL/6J and SJL/J genetic backgrounds through oral glucose tolerance testing at 4, 5, 6, and 8 weeks of age to evaluate glucose tolerance and glycemic trajectory. All F2 mice were genetically monitored for single nucleotide polymorphisms (SNPs) between the C57BL/6J and SJL/J backgrounds, allowing us to identify modifying loci on 6 chromosomes (2, 126 Mbp; 3, 59 Mbp and 103 Mbp; 7, 47 Mbp; 11 45 Mbp; 13, 4 Mbp; and 16, 85-97 Mbp), containing 172 candidate genes. For example, known MODY genes *Abcc8* (Sur1) and *Kcnj11* (Kir6.2) are within the QTL region on chromosome 7 and contain polymorphisms between backgrounds that may impart combinatorial effects on the phenotypic penetrance of MafA^S64F^-mediated dysglycemia.

## INTRODUCTION

Although rare, diabetes caused by single gene variants (monogenic diabetes) has led to the identification of essential regulators of glycemia and expanded our understanding of the pathophysiology of polygenic diabetes (e.g., T2D)^1,2^. Many of the genes implicated in monogenic diabetes comprise transcription factors (TFs) enriched in pancreatic islet cells, such as the large V-Maf Avian <u>m</u>usculo<u>a</u>poneurotic fibrosarcoma (Maf<u>A</u>) TF. Notably, MafA is enriched in the pancreatic β-cells^3–7^, promoting β-cell differentiation and transcriptionally regulating genes involved in nearly every step of glucose-stimulated insulin secretion (GSIS), a defining functional feature of mature islet β-cells^8–11^. Genetic mouse models harboring a β-cell-specific deletion of *Mafa* show dysglycemia^12^, while exogenous expression of *Mafa* increases GSIS in normally non-glucose responsive MafA^Low^ neonatal rat islets^13^. Further, MafA facilitates β-cell maturation and GSIS in mouse and human β-like stem-cells^13–16^. These results collectively demonstrate MafA is essential for postnatal β-cell function.

Clinically, human MafA is one of the first factors downregulated in T2D β-cells^17^. Moreover, a MafA missense variant (MafA^Ser64Phe,^ or MafA^S64F^) predisposes adult human carriers to disease in adulthood (late 30’s): either maturity-onset diabetes of the young (β-cell dysfunction; MODY) or non-syndromic insulinomatosis (functional β-cell tumors)^18^. Intriguingly, male carriers are more prone to diabetes, while female carriers have a greater susceptibility to insulinomatosis resulting in hypoglycemia^18^. A second MafA missense variant (MafA^Thr57Arg^) was independently identified with a similar sex-biased phenotype^19^.

Our group has developed a mouse strain harboring the S64F mutation in the endogenous *Mafa* gene that demonstrated the expected sex-dependent phenotypes of humans, with males having impaired glucose tolerance and insulin secretion and females having an increased insulin secretory response^20^. Peripheral insulin sensitivity was not grossly affected^20^, which revealed β-cell-intrinsic defects in the phenotypes. We have recently shown the ability of genetic background to modulate the glycemic phenotype of the *Mafa*^S64F/+^ mice^21^, leading us to investigate genetic modifiers of MafA^S64F^ in the present study. We generated *Mafa*^S64F/+^ F2 mice derived from C57BL/6J (C57) and SJL/J (SJL) backgrounds and carried out oral glucose tolerance testing (GTT). Results of quantitative trait loci (QTL) mapping of these data were overlaid with MafA CUT&RUN and RNA sequencing datasets^21^ to identify candidate genes influencing Mafa^S64F/+^ glycemia.

## METHODS

### Mouse husbandry

All animal studies were reviewed and approved by the Vanderbilt University Institutional Animal Care and Use Committee. Mice were housed in conventional cages (2-5 mice/cage) at 22°C using a 12h light cycle starting at 0700h, with ad libitum access to water and standard chow diet. Mice were cared for according to the Vanderbilt University Department of Animal Care and the Institutional Animal Care and Use Committee of Animal Welfare Assurance Standards and Guidelines.

### Mice Line Generation

The MafA^S64F^ mouse line was originally generated as described previously^20^, and the C57BL/6J MafA^S64F^ line was generated by backcrossing the MafA^S64F^ mutation into the C57BL/6J (Jackson Labs) mouse line as previously discussed^21^. These mice were bred a single time with the SJL/J (Taconic) to generate the CHA F1 line with an approximately 50:50 contribution from both genetic backgrounds. A Mafa^S64F/+^ CHA F1 male was bred with a wild-type C57BL/6J or SJL/J female to generate the C57 N1 and SJL N1 lines respectively (**Figure 1A**). *Mafa^S^*^64F/+^ CHA F1s were bred with *Mafa*^WT^ CHA F1s to generate CHA F2 progeny. Genetic background monitoring was performed by Transnetyx using the Mini Mouse Universal Genotyping Array (MiniMUGA)^22^.

**Figure 1.**
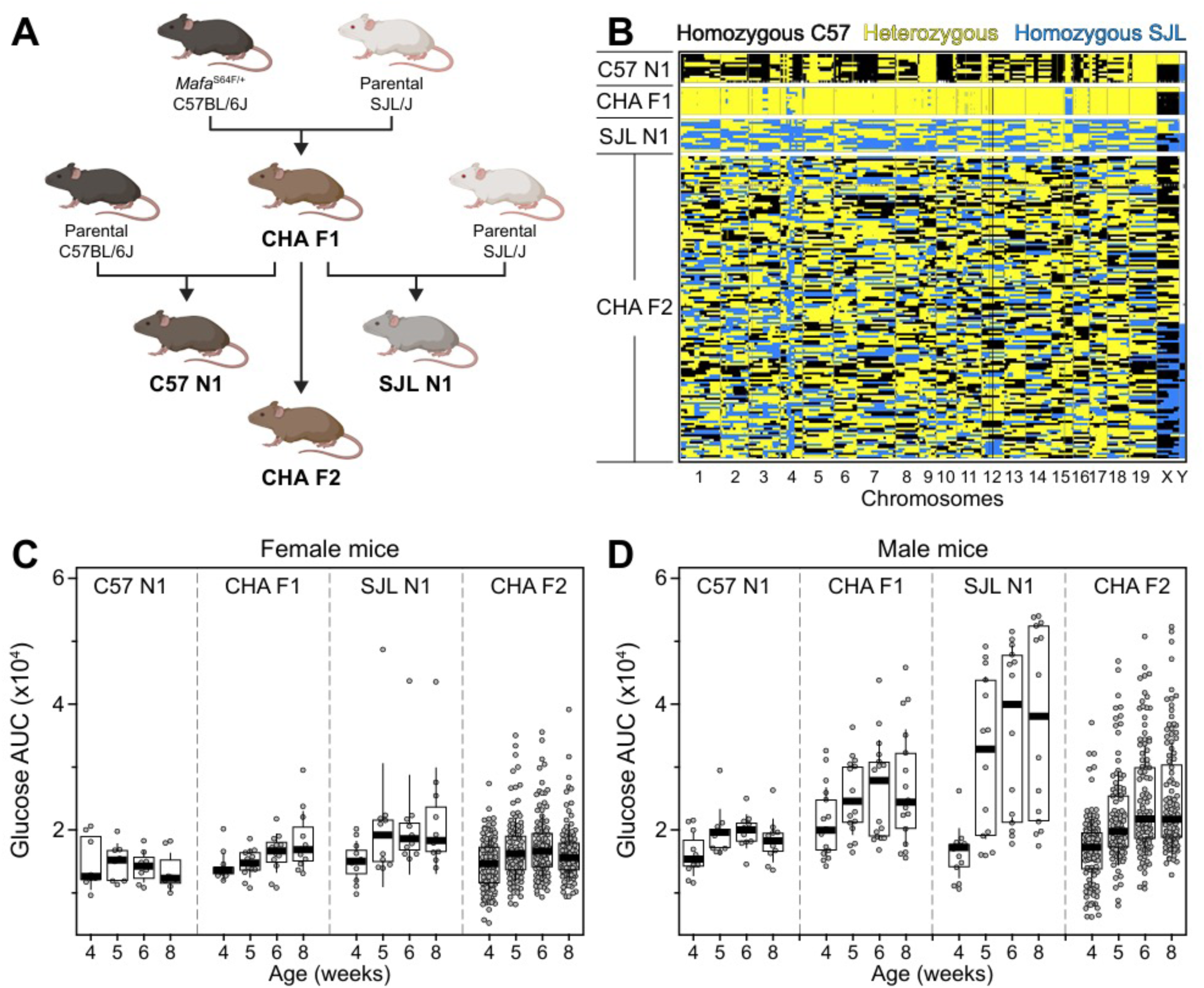
Genetic background modulates phenotype of both male and female *Mafa*^S64F/+^ mice. **A.** Breeding strategy used to generate the four mouse lines used for these experiments. **B.** Genetic monitoring results of all *Mafa*^S64F/+^ mice. **C.** Area under the curve (AUC) of oral glucose tolerance testing of female *Mafa*^S64F/+^ mice. Data is represented as mean +/- 1 SD. C57 N1 n = 5, CHA F1 n = 8, SJL N1 n = 6, CHA F2 n = 73. **D.** AUC of oral glucose tolerance testing of male *Mafa*^S64F/+^ mice. Data is represented as mean +/- 1 SD. C57 N1 n = 3, CHA F1 n = 10, SJL N1 n = 8, CHA F2 n = 61.

### Oral glucose tolerance testing

D-glucose was prepared in sterile PBS (20% w/v). Glucose tolerance testing was performed on all mice in the afternoon by giving a bolus of D-glucose (2 mg/g body weight) through oral gavage after a 5-hour fast. Blood glucose was measured using a FreeStyle glucometer (Abbott Diabetes Care) before the bolus (0 minutes) and at 15, 30, 60, and 90 minutes after the bolus. Glucose tolerance testing was done on mice at 4, 5, 6, and 8 weeks of age.

### SNP identification

We used the single nucleotide polymorphism (SNP) search tool at MGI (https://www.informatics.jax.org/snp)^23^ to identify all SNPs that are polymorphic between C57BL/6J and SJL/J. To annotate these SNPs and their predicted consequence we utilized the Variant Effect Predictor (VEP) tool (v114.0)^24^, which allowed us to recover several variant classes associated with coding and non-coding genomic regions. We further leveraged comprehensive annotations available from the Ensembl^25^ and GENCODE^26^ databases to predict the functional consequences of the annotated SNPs. SNPs that were annotated as resulting in high (e.g., STOP gained, splice junction) or moderate (e.g., missense) impact consequences were further characterized by their relative conservation.

To quantify evolutionary constraint at single base-pair resolution, sequence conservation scores (phastCons scores) were extracted using the phastCons35way.UCSC.mm39 annotation package^27^ implemented within the GenomicScores^28^ framework in R. PhastCons scores evaluate the posterior probability—ranging from 0.0 (unconstrained) to 1.0 (highly conserved)—that a specific nucleotide belongs to a conserved genomic element, derived from a phylogenetic Hidden Markov Model (HMM) spanning a 35-way vertebrate species alignment against the GRCm39/mm39 mouse reference genome. These conservation scores were integrated with predicted functional consequence severity (VEP) and amino acid tolerance predictions from SIFT^29^ to establish a composite variant priority score for candidate causal QTL discovery. This allowed us to identify 62 high-impact variants and 4,496 moderate-impact variants between C57BL/6J and SJL/J mice with a phastConservation score > 0.6.

### QTL mapping

Genetic analysis was performed using quantitative trait loci (QTL) methods implemented in R/qtl^30,31^. Because MafA^S64F^ is known to have sex-dependent effects on glycemia, sex was used as an additive covariate in QTL-mapping. This involved genome scans of GTT measurements, including summary traits based on AUC (area under the curve) and further meta-traits over the multiple weeks of assays. Peaks above a permutation threshold (∼4) and 95% approximate confidence intervals were reported.

### MafA RNA sequencing

Bulk RNA sequencing data and methods have previously been published.^21^ In short, bulk RNA sequencing was performed on islets isolated from 5-6-week old males of both CHA F1 and fully backcrossed C57 (Transnetyx Genetic Monitoring, >95% C57BL/6J) mice. Samples with RIN > 8 were sequenced on an Illumina NovaSeq6000, and data was processed and interpreted using Genialis bioinformatics software. DESeq2 was used for differential gene expression analysis and statistical comparison.

### MafA CUT&RUN sequencing

CUT&RUN data and methods have previously been published^21^. In short, CUT&RUN was performed on 500,000 dispersed mouse MIN6 cell nuclei using the CUTANA ChIC/CUT&RUN protocol v3.1 (EpiCypher). DNA libraries were sequenced as PE150 reads on the NovaSeq platform. The pipeline performs adaptor trimming, alignment to the GRCm38 reference genome, filtering against the mm10 blacklist regions, spike-in normalization and peak calling using MACS2. Quality was confirmed by fragment length distribution and fingerprint plots. Nearest gene analysis was performed with peakScout^32^. Coverage plots were created using ggcoverage^33^.

### Integration of human GWAS

To assess the whether a gene is associated with glycemic traits in human GWAS, we used Human Genetic Evidence (HuGE) scores, which combines common variant data from GWAS with rare variant data from whole-exome sequencing, providing a gene-centric score of trait associations^34,35^.

### MafA Discovery App development

MafA CUT&RUN binding peaks (∼33,000) were filtered to just those containing one or more genetic variants (range 1 to 66, median = 2) between C57BL/6J and SJL/J to yield ∼10,500 SNP-containing peaks. We performed RNA sequencing of pancreatic islets from *Mafa*^S64F/+^ CHA F1 mice that were backcrossed to either C57 or SJL mice, enabling us to identify ∼4,400 differentially expressed genes (DEGs, log_2_FC >1) that distributed in the following manner: 1,069 specific to C57; 2,334 specific to SJL; 998 that were DE and in the same direction for both C57 and SJL; and 7 that were DE but in the opposite direction. The MafA Discovery App (https://chalab2022.github.io/MafADiscovery/) merges these data sets (SNP-containing MafA peaks, DEGs and high/moderate impact SNPs) to allow us to identify causal candidate genes that occur within the 95% confidence intervals (CI) of the glycemic-associated QTL identified in the F2 mice.

## RESULTS

### Mouse genetic background significantly affects penetrance of MafA-MODY

A mouse model of the MafA^S64F^ mutation mirrors several aspects of the clinical phenotype, with male mouse carriers developing hyperglycemia and decreased glucose tolerance while female mice show hypoglycemia^20^. We have previously shown the relative proportions of inbred genetic backgrounds can modulate the phenotypic penetrance of male *Mafa*^S64F/+^ mice, but such marked differences in penetrance were not seen in female *Mafa*^S64F/+^ mice^21^. In addition to the protective effect of the C57BL/6J “C57” background seen previously, we have since identified a diabetogenic effect from the SJL/J “SJL” background (**Supplemental Figure 1A-C**). These results show that even one generation of backcross permits clear detection of disparate glycemic trends between strains in male mice and led us to investigate the genomic differences between C57 and SJL parental strains in males.

Comparing unique SNPs within parental strains did not identify high impact variants in MafA or known MafA interacting proteins

To nominate potential genetic modifiers between C57 and SJL mice which may influence phenotypic penetrance in MafA^S64F^ males, we identified all single nucleotide polymorphisms (SNPs) and insertions/deletions (indels) between C57 and SJL parental strains. We extracted genotype calls for these variants from the latest release of the Mouse Genomes Project, a resource containing sequencing information for 52 mouse strains, enabling us to identify ∼5.3 million high-confidence variants between C57 and SJL. We further leveraged comprehensive annotations available from the Ensembl^25^ and GENCODE^26^ databases to predict the functional consequences of these variants, which allowed us to focus on those that yield a high (e.g., stop gained) or moderate (e.g., missense variant) impact on protein-coding genes. This filtering strategy produced a list of 343 and 14,401 high and moderate impact variants in 311 and 4,987 genes, respectively (**Supplemental Table 1**). No SNPs were identified within known MafA interacting proteins^7,36^ or within MafA itself, so we decided to pursue an unbiased investigation of Mafa^S64F/+^ mice to elucidate novel genetic modifiers causing the observed phenotypic differences.

### QTL-mapping identifies 6 significant genomic loci affecting MafA^S64F^-mediated dysglycemia

Exploiting these differences in glucose tolerance between genetic backgrounds, we generated an F2 C57xSJL intercross of *Mafa*^S64F/+^ mice (CHA F2; **Figure 1A**) to map novel quantitative trait loci (QTL) modulating MafA^S64F^. Genetic monitoring results confirmed increased recombination events and genetic variability of the CHA F2 generation when compared to the backcrossed (C57 N1 and SJL N1) mice and the CHA F1s (**Figure 1B**). Oral glucose tolerance testing (GTT) was performed on backcrossed and intercrossed (CHA F1 and CHA F2) mice at 4, 5, 6, and 8 weeks of age to longitudinally capture glucose tolerance and trajectory (**Supplemental Figure 1A-D**). As we had previously observed^21^, only mild effects from age and genetic background were observed on GTT area under the curve (AUC) for female *Mafa*^S64F/+^ mice (**Figure 1C**), while more dramatic effects from both age and genetic background were observed for male *Mafa*^S64F/+^ mice (**Figure 1D**). As predicted, the range of GTT AUCs observed in the F2s fully encompassed the range bounded by those observed in the backcrossed mice (**Figure 1C-D**).

We performed QTL analysis of the direct glycemic traits (AUC at 4, 5, 6, and 8 weeks), as well as longitudinal glycemic meta-traits (AUC of AUCs, slope of the AUCs, and AUC_8-week_ –AUC_4-week_), with sex as an additive covariate. Significant loci, as defined by an LOD score above 4.03, were identified on five autosomal chromosomes: 3, 7, 11, 13, and 16 (**Figure 2A**, **Table 1**). Additionally, a suggestive locus was identified at 126 Mbp on chromosome 2. At all identified loci, SJL alleles had pro-diabetic effects on their associated traits (**Figure 2B**): the lowest glucose readings were observed in mice which had homozygous C57 alleles at that locus and these effects progressively increased with heterozygous C57/SJL and homozygous SJL mice. The effects on 8-week AUC on chromosome 2 and 5-week AUC on chromosome 7 were generally unchanged between heterozygous C57/SJL compared to homozygous SJL. Both 5-week AUC on chromosome 13 and 4-week AUC on chromosome 11 had worsened effects when heterozygous at those loci compared to being homozygous for either background.

**Figure 2.**
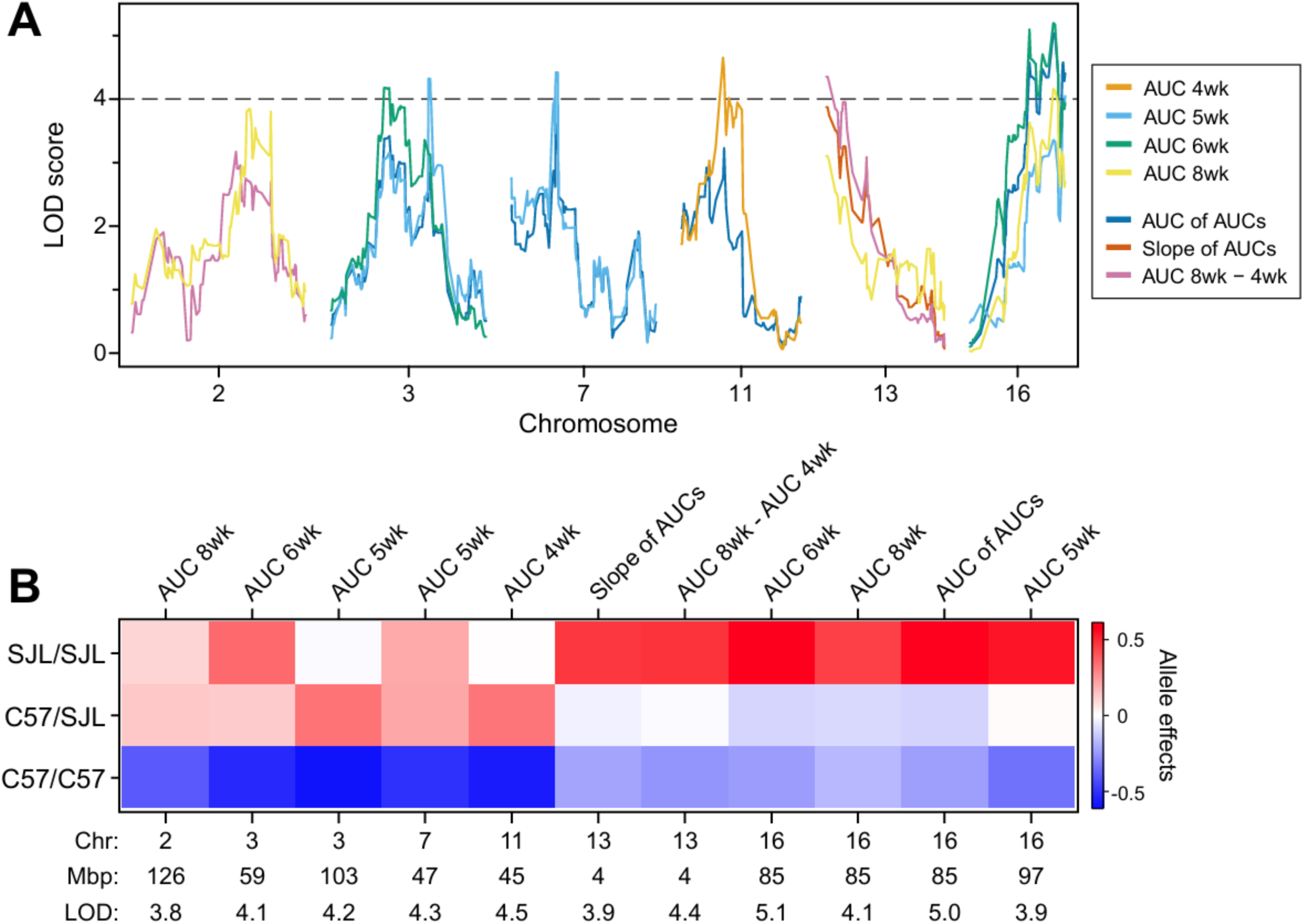
SJL alleles on multiple chromosomes worsen AUC phenotype. **A.** Genome scans mapped against LOD scores of glycemic traits (AUC at 4, 5, 6, and 8 weeks) and trajectory of glycemia (AUC of AUCs, Slope of AUCs, and AUC4week – AUC8week) from CHA F2 mice. Significant QTLs identified on chromosomes 3, 7, 11, 13, and 16. **B**. Allele effects at significant loci for the corresponding glycemia or trajectory trait. AUC, area under the curve of glucose tolerance tests. LOD, logarithm of the odds.

**Table 1.**
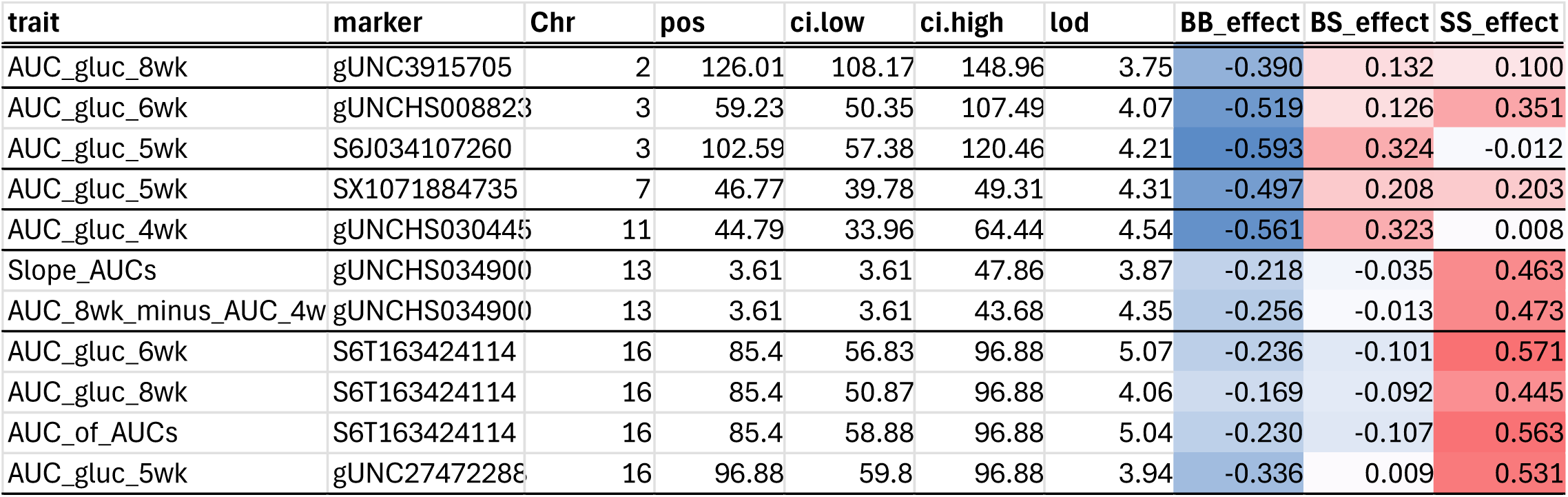
QTLs identified for CHA F2 mice based on direct glycemic traits or meta-glycemic traits.

| trait | marker | Chr | pos | ci.low | ci.high | lod | BB_effect | BS_effect | SS_effect |
|---|---|---|---|---|---|---|---|---|---|
| AUC_gluc_8wk | gUNC3915705 | 2 | 126.01 | 108.17 | 148.96 | 3.75 | -0.390 | 0.132 | 0.100 |
| AUC_gluc_6wk | gUNCHS008823 | 3 | 59.23 | 50.35 | 107.49 | 4.07 | -0.519 | 0.126 | 0.351 |
| AUC_gluc_5wk | S6J034107260 | 3 | 102.59 | 57.38 | 120.46 | 4.21 | -0.593 | 0.324 | -0.012 |
| AUC_gluc_5wk | SX1071884735 | 7 | 46.77 | 39.78 | 49.31 | 4.31 | -0.497 | 0.208 | 0.203 |
| AUC_gluc_4wk | gUNCHS030445 | 11 | 44.79 | 33.96 | 64.44 | 4.54 | -0.561 | 0.323 | 0.008 |
| Slope_AUCs | gUNCHS034900 | 13 | 3.61 | 3.61 | 47.86 | 3.87 | -0.218 | -0.035 | 0.463 |
| AUC_8wk_minus_AUC_4w | gUNCHS034900 | 13 | 3.61 | 3.61 | 43.68 | 4.35 | -0.256 | -0.013 | 0.473 |
| AUC_gluc_6wk | S6T163424114 | 16 | 85.4 | 56.83 | 96.88 | 5.07 | -0.236 | -0.101 | 0.571 |
| AUC_gluc_8wk | S6T163424114 | 16 | 85.4 | 50.87 | 96.88 | 4.06 | -0.169 | -0.092 | 0.445 |
| AUC_of_AUCs | S6T163424114 | 16 | 85.4 | 58.88 | 96.88 | 5.04 | -0.230 | -0.107 | 0.563 |
| AUC_gluc_5wk | gUNC27472288 | 16 | 96.88 | 59.8 | 96.88 | 3.94 | -0.336 | 0.009 | 0.531 |

### MafA Discovery App helps to identify candidate genes

We developed a discovery application to help narrow down potential modifying candidate genes of interest within QTL regions (**Figure 3A**). The MafA Discovery App (https://Chalab2022.github.io/MafADiscovery) incorporates our previously published bulk RNA sequencing and MafA CUT&RUN data^21^ to visually overlay DEGs uniquely changed in each genetic background (i.e., C57 and CHA F1) and known genomic regions of MafA occupancy by CUT&RUN. We also overlaid coding variant SNPs unique between the C57 and SJL parental backgrounds (**Table 1**) with their phastCons score and relative impact. Finally, genes that fit these criteria were further investigated for their effects on human glycemia from GWAS studies using a HuGE score and their possible relevance to MafA in the literature.

**Figure 3.**
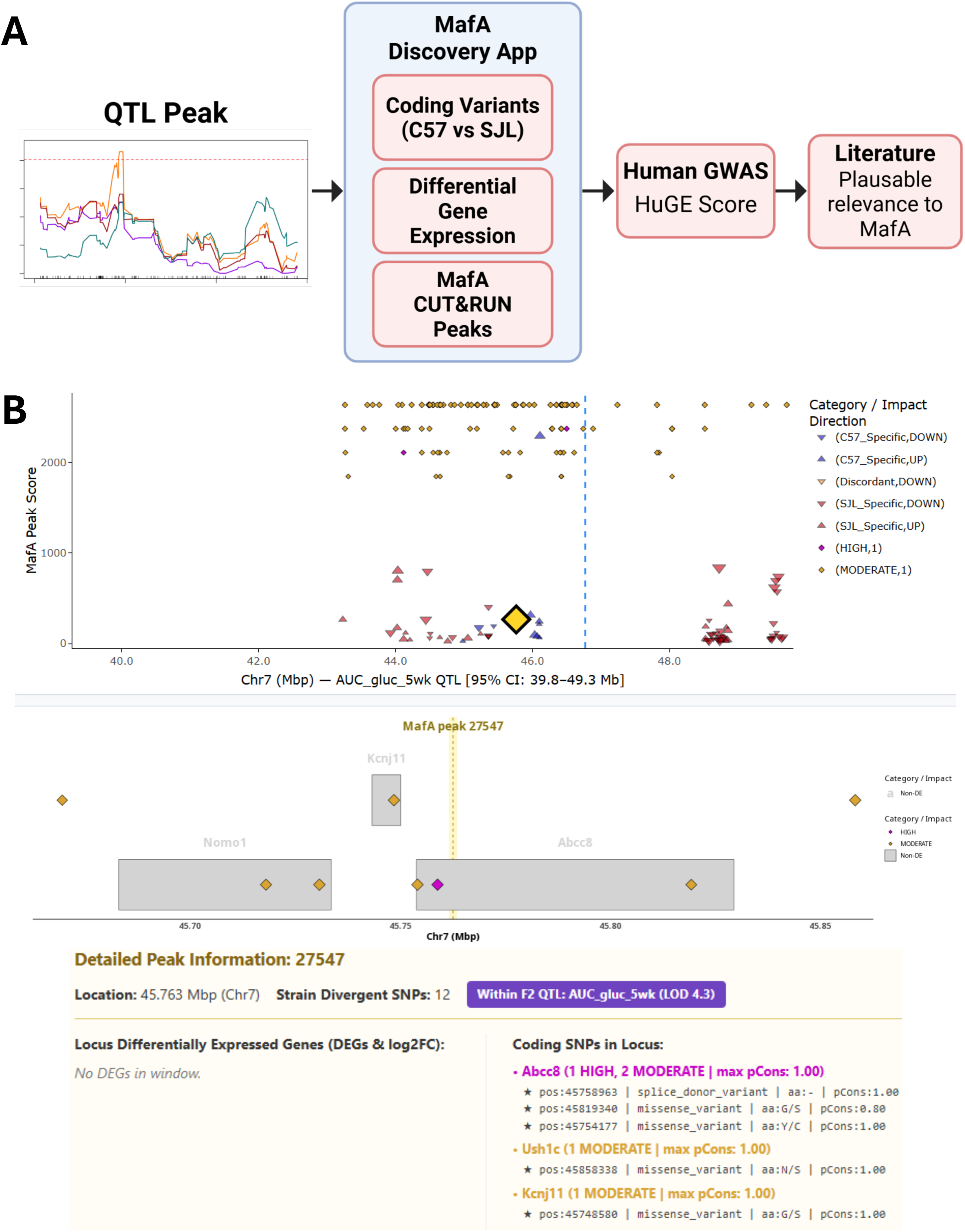
Prioritization of genes identifies *Abcc8* and *Kcnj11* as potential candidate genes. **A.** Pipeline for candidate gene prioritization within QTL regions. Genes within the confidence interval are further inspected for coding variants between the C57 and SJL backgrounds, with priority to those with a phastCons score > 0.6, bulk RNA-seq differential expression, and proximity to a peak from MafA CUT&RUN^21^. These three factors can be visualized in the MafA Discovery App. Genes are further prioritized by their HuGE score for glycemic association in human GWAS^34^. Finally, literature searches for a plausible relevance to MafA were performed. **B.** Screenshot of the MafA Discovery App showing the significant QTL on chromosome 7. The top graph shows the 95% confidence interval locus, and the peak is indicated by a blue dotted line. DEGs (triangles) and coding variants (diamonds) within that region are shown. The lower graph is a zoomed-in view focused on *Abcc8* and *Kcnj11*, with diamonds indicating coding variants. The nearest MafA CUT&RUN peak to the selected gene is indicated by the yellow dotted line. Detailed information of genes within the view is below. Abcc8 has 2 moderate and 1 high impact SNPs between C57 and SJL, while *Kcnj11* has one moderate SNP.

Using this pipeline, we identified 34 coding genes on chromosome 2, 39 on chromosome 3, 21 on chromosome 7, 31 on chromosome 11, 28 on chromosome 13, and 19 on chromosome 16 (**Supplemental Table 2**). For example, among the genes encoded within the chromosome 7 QTL are *Abcc8* and *Kcnj11*, known MODY genes^37,38^. *Abcc8* contains three SNPs with phastCons scores greater than 0.6, one high impact splice donor variant and two moderate impact missense mutations, and *Kcnj11* has one moderate impact SNP with a phastCons score of 1 (**Figure 3B**). Additionally, *Ucn3* in the chromosome 13 QTL is a marker of β-cell maturity and function^39^. *Ucn3* is down-regulated in CHA F1 *Mafa*^S64F/+^ male mice compared to wild-type littermates (log2FC = -1.59), but not in C57 *Mafa*^S64F/+^ male mice (**Figure 4**). The combinatorial impact of these critical genes likely contributes to the phenotypic differences between genetic backgrounds. Additionally, 15 genes were identified to have both SNPs between C57 and SJL and differential expression between *Mafa*^S64F/+^ and WT male mice of either background, as well as a HuGE score >1 on one or more glycemic traits (**Table 2**).

**Figure 4.**
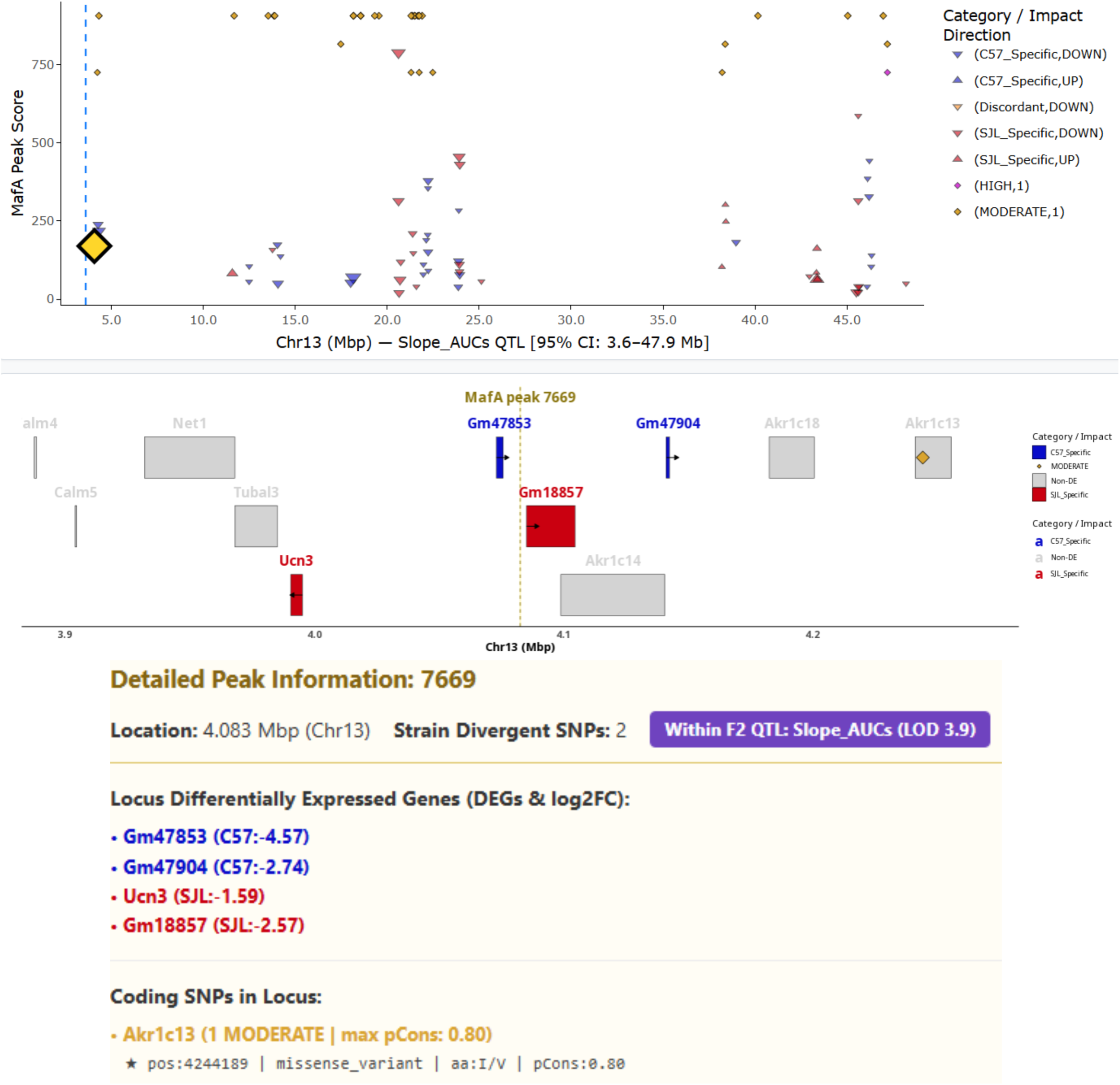
Candidate prioritization highlights *Ucn3* as a potential MafA^S64F^ modifier. **A.** Screenshot of the MafA Discovery App showing the significant QTL on chromosome 13. The top graph shows the 95% confidence interval locus, and the peak is indicated by a blue dotted line. All the differentially expressed genes (triangles) and coding variants (diamonds) within that region are shown. The lower graph is zoomed-in view focused on *Ucn3*, colored red to indicate differential expression in the F1 mice. The nearest MafA CUT&RUN peak to the selected gene is indicated by the yellow dotted line. Detailed information of genes within the view is below. *Ucn3* is downregulated in CHA F1 *Mafa*^S64F/+^ male mice compared to WT littermates, with a log2FC of -1.59.

**Table 2.** Genes within QTLs containing highly conserved SNPs between C57 and SJL as well as differential expression between *Mafa*^S64F/+^ and WT male mice.

| Gene | Chr | Start Mbp | End Mbp | Total Conserved SNPs | Highest phastCons | C57 log2FC | F1 log2FC |
|---|---|---|---|---|---|---|---|
| ABI3BP | 16 | 56.29824 | 56.5105 | 5 | 1 |  | **1.416** |
| ADAMTS5 | 16 | 85.65306 | 85.69872 | 2 | 1 |  | **1.413** |
| BCHE | 3 | 73.54314 | 73.61575 | 1 | 1 | **1.392** | **1.243** |
| DLL4 | 2 | 119.1563 | 119.1664 | 1 | 0.9 | **-1.126** |  |
| DPYD | 3 | 118.3558 | 119.2266 | 1 | 1 |  | **1.165** |
| FBN1 | 2 | 125.1425 | 125.3499 | 1 | 1 |  | **1.38** |
| FSTL5 | 3 | 75.98158 | 76.61733 | 2 | 1 | **1.102** | **1.502** |
| HS3ST3B1 | 11 | 63.77662 | 63.81312 | 1 | 1 | **1.438** | **1.195** |
| Nav2 | 7 | 48.55846 | 49.25984 | 1 | 1 |  | **-1.199** |
| NOTCH2 | 3 | 97.92084 | 98.05768 | 1 | 0.8 |  | **1.247** |
| PTTG1 | 11 | 43.31108 | 43.31708 | 1 | 1 |  | **-1.653** |
| RALGAPA2 | 2 | 146.0818 | 146.3543 | 6 | 1 |  | **1.15** |
| RYR2 | 13 | 11.56799 | 12.12183 | 1 | 1 |  | **2.444** |
| SYT6 | 3 | 103.4825 | 103.5529 | 1 | 0.9 |  | **1.454** |
| TRIM58 | 11 | 58.53129 | 58.54323 | 1 | 0.9 | **5.092** |  |

## DISCUSSION

Common genetic variants have been identified to influence monogenic HNF-MODY^40^ and GCK-MODY^41^ penetrance, demonstrating the presence of modifying genetic factors on clinical disease onset and severity. Moreover, several MODY genes are implicated in broader type 2 diabetes^1^, suggesting that the intersection of MODY and modifying genes may also impact the risk and predisposition for polygenic type 2 diabetes. We recently identified that genetic background has a profound impact on the male diabetes penetrance and islet aging on a mouse model harboring a human MafA-MODY variant^21^. Using controlled breeding strategies to change genetic background contributions, we “tuned” diabetes penetrance (i.e., prevented or worsened) in this mouse model to investigate the influence of genetic landscape and modifiers on β-cell function in response to a pathogenic MODY variant. However, specific genetic loci that contributed to the dramatic differences in *Mafa*^S64F/+^ male phenotypes between genetic backgrounds were not resolved.

Here we constructed an F2 cross between C57BL/6J and SJL/J Mafa^S64F/+^ mice to map disease-modifying loci. We identified QTLs at chromosomes 2 (126 Mbp), 3 (59 Mbp and 103 Mbp), 7 (47 Mbp), 11 (45 Mbp), 13 (4 Mbp), and 16 (85-97 Mbp). We referenced SNPs in protein-coding genes within these candidate loci against previously published MafA RNA sequencing and CUT&RUN datasets, from which we constructed a MafA Discovery App for public use. This has led to the identification of MafA-regulated candidates such as Abcc8, Kcnj11, and Ucn3 for further validation.

Together, *Abcc8* (encoding SUR1) and *Kcnj11* (encoding Kir6.2) form the ATP-sensitive potassium (K_ATP_) channel on β-cells to help with membrane depolarization mediating glucose-stimulated insulin secretion^42^. Genetic variants of both genes have been linked to multiple diseases, with loss-of-function mutations leading to congenital hyperinsulinism (persistent hypoglycemia)^42,43^, and gain-of-function mutations causing neonatal diabetes^43,44^ or MODY^37,38^. Mice with a dominant negative mutation of *Kcnj11* show improved neonatal blood glucose levels, but elevated glucose levels by 4-weeks^45^, similar to the phenotypic transition seen in male *Mafa^S64F/+^* mice with a mixed genetic background^20^. SJL mice have two missense mutations and one splice donor variant in *Abcc8*, as well as one missense variant in *Kcnj11*, thus, one or more of these mutations could be influencing the function of the β-cell K_ATP_ channels and thus modulating the pathogenic effects of MafA^S64F^.

*Ucn3* (encoding urocortin 3, Ucn3) is a β-cell maturity marker, appearing later in β-cell development^46^. Ucn3 is co-released with insulin by β-cells, and binds to receptors on δ-cells to potentiate somatostatin release^46^. The downregulation of *Ucn3* in *Mafa*^S64F/+^ male CHA F1 mice, but not C57 mice, correlates with worsened glucose tolerance, pointing towards a potential role of β-cell maturity and cross-talk with δ-cells in glycemic phenotype of *Mafa*^S64F/+^ mice. In sum, this study identifies the influence of genetic background and ancestry in glycemia and, in this model, identifies combinatorial impact of critical β-cell enriched genes.

As we previously reported, MafA-MODY is clinically rare. Only 3 families have been identified thus far to carry this mutation (MafA^S64F^) with sex-dependent phenotypes^18,47^, while another missense mutation (MafA^Thr57Arg^) was identified in one family^19^. No human carriers of the MafA^S64F^ mutation were identified in the gnomAD (https://gnomad.broadinstitute.org/), Ensembl (https://useast.ensembl.org/Homo_sapiens/Info/Index) and GWAS catalogue (https://www.ebi.ac.uk/gwas/home). However, heterozygous impairment of causative MODY genes also overlap with T2D^1,2^, suggesting mechanistic overlap between T2D and monogenic diabetes. Indeed, MafA has a profound role in β-cell biology, and efforts to mechanistically investigate the influence of a single gene variant is not only far simpler to unraveling the regulation of polygenic diabetes but can also significantly improve our understanding of the complexity of T2D through shared mechanisms.

We acknowledge limitations to this study. We only investigate the physiologic readout of glucose tolerance testing without other physiologic measurements taken such as insulin secretion or MafA protein levels. We have measured insulin and C-peptide secretion in *Mafa*^S64F/+^ mice from both the F1 mixed background and a fully backcrossed C57 background, and those values corresponds well to GTT curves^21^, suggesting GTT as an potential proxy for insulin secretion in MafA^S64F^ islets. Further, the F2 generation yielded relatively large genomic loci harboring many candidate genes. Future studies with further intercrossing can be pursued to validate the candidates identified presently. Additionally, identified candidates need to be validated with causative molecular investigation. Multiple candidates may act in concert and may require multiplexed validation.

Our study demonstrates the functionality of identifying MafA-regulated genetic modifiers of glycemia using a robust mouse line modelling MafA-MODY with adequate signal:noise in glycemic traits between strains. Further analysis of candidate modifiers conserved in humans can improve strategies for diabetes risk stratification and therapies to optimize β-cell function in people at risk of diabetes.

## Article Contributions

The study was conceived and designed by MM, MK, and JC . Funding was obtained by JC. All authors contributed to the acquisition, analysis or interpretation of the data. Data and statistical analyses were performed by MM, LJ, DL, BY, MK and JC. MM, BY, MK, and JC drafted the manuscript, and all authors critically revised it for important intellectual content. MK and JC supervised the work. All authors approved the final version of the manuscript. JC takes responsibility for the integrity of the work as a whole.

## SUPPLEMENTAL ITEMS

**Supplemental Figure 1.**
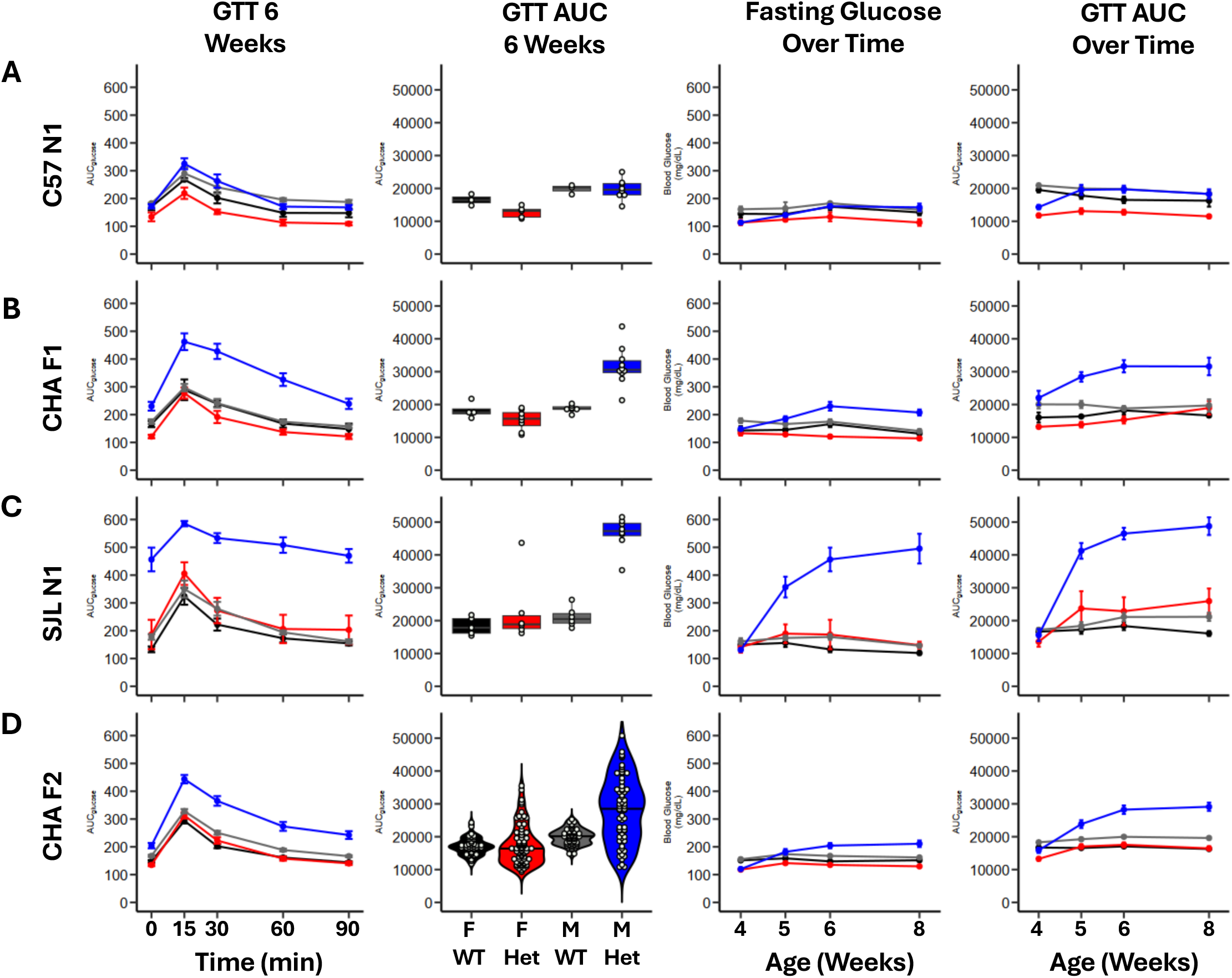
Glucose tolerance testing across mouse generations. A. Glycemic profile of C57 N1 mice between 4 and 8 weeks, including the GTT and GTT AUC at 6 weeks, fasting glucose levels over time, and GTT AUC over time. B. Glycemic profile of CHA F1 mice between 4 and 8 weeks. C. Glycemic profile of SJL N1 mice between 4 and 8 weeks. D. Glycemic profile of CHA F2 mice between 4 and 8 weeks. Black, Female WT. Red, Female *Mafa*^S64F/+^. Gray, Male WT. Blue, Male *Mafa*^S64F/+^. GTT, glucose tolerance test. AUC, area under the curve.

**Supplemental Table 1. Differential SNPS between SJL and C57 genetic background mice predicted to be missense mutations.**

**Supplemental Table 2. Prioritization of genes within QTLs using SNPs, differential gene expression and HuGE scores.**

